# Three-dimensional humanized gingival tissue model to study oral microbiome

**DOI:** 10.1101/2022.07.17.500348

**Authors:** Miryam Adelfio, Zaira Martin-Moldes, Joshua Erndt-Marino, Lorenzo Tozzi, Margaret J. Duncan, Hatice Hasturk, David L. Kaplan, Chiara E. Ghezzi

**Affiliations:** Department of Biomedical Engineering, University of Massachusetts Lowell, Lowell, Massachusetts, USA; Department of Biomedical Engineering, Tufts University, Medford, Massachusetts, USA; Department of Microbiology, The Forsyth Institute, Cambridge, Massachusetts, USA; Center for Clinical and Translational Research, The Forsyth Institute, Cambridge, Massachusetts, USA

**Keywords:** tissue model, host-pathogen interactions, microbiome, eubiosis, gingiva, periodontitis, silk

## Abstract

The oral cavity contains different microenvironments, as the non-shedding surface of the teeth and the epithelial mucosa, where oral barriers and microbial communities coexist. The interactions and balances between these two communities are responsible for oral tissue homeostasis or dysbiosis, that ultimately dictate health or disease. Disruption of this equilibrium is the first necessary step towards chronic inflammation and permanent tissue damage in the case of chronic periodontitis. There are currently no experimental models able to mimic the structural, physical, and metabolic conditions present in the oral gingival tissue to support the long-term investigation of host-pathogens unbalances. Herein, we report a 3D anatomical gingival *in vitro* model based on human primary culture that recapitulates the native tissue organization, and a native oxygen gradient within the gingival pocket to support human microbiome persistence with a physiologically relevant level of microbial diversity as well as native spatial organization. The modulation of inflammatory markers in the presence of oral microbiome suggested the humanized functional response of this model. The model will be used in future studies to investigate host-pathogen unbalances in gingivitis and periodontal disease.

## 1 Introduction

The dynamic and polymicrobial oral microbiome is the initiator of diseases such as periodontitis and dental caries, globally two of the most predominant microbially induced disorders ^[1]^. The oral cavity contains different microenvironments, that colonize the non-shedding surface of the teeth and the epithelial mucosa, where oral barriers and microbial communities coexist ^[2]^. The interactions and balances between these two communities are responsible for oral tissue homeostasis or dysbiosis, that ultimately dictate health or diseased tissue states, respectively. Disruption of this equilibrium is the first necessary step that ultimately leads to chronic inflammation and permanent tissue damage in the case of periodontitis ^[3]^. Due to mild initial symptoms as well as the large variety of its clinical manifestations, the overall model of disease initiation and progression are difficult to establish ^[4]^. While gingival health and gingivitis have many clinical features, case definitions are primarily predicated on presence or absence of bleeding on probing ^[2]^. In fact, the current working polymicrobial synergy and dysbiosis (PSD) model suggests that the disease is not originated by individual causative periopathogens (*i*.*e*., red complex), but by continuous cyclic interactions between physically and metabolically integrated polymicrobial communities and an unbalanced host inflammatory response ^[5] [4] [2]^. Thus, emphasis is placed on identifying investigative tools to systematically study these complex interactions, rather than scrutinizing individual pathogenic elements, as in the past conventional culture-based approaches ^[6] [7] [8]^.

Host-pathogen interactions have been mainly investigated in clinical and animal models, but also in *in vitro* studies. Considering the interindividual variability and diversity shown in clinical investigations ^[9]^, animal and *in vitro* models have offered a great advantage in elucidating the trajectory of the disease. Humanized mice can, in fact, provide significance to clinical observations, via mechanically or bacterial inoculum-induced periodontitis models ^[10] [11]^; however, although they are considered a means of translational applicability in periodontal disease, they do not reflect the complex pathological scenario shown in the human condition due to differences in microbiome composition and induction of the dysbiotic condition (*i*.*e*., ligature) ^[12]^. Current *in vitro* strategies are limited to two-dimensional culture systems based on immortalized cells that are only functional for a short window of time (maximum 7 days) ^[13] [14] [15]^. Such systems have been used to test irritant responses of new dental materials, dentifrices, and oral care consumer products, but are unable to maintain the complexity of the oral pathogen community organization, due to the lack of the native oxygen and metabolic conditions. Moreover, current experimental models to study host-pathogens interplays rely on the use of planktonic bacteria cultures that neglect the interspecies interactions required for attachment, colonization, and regulation of host mucosal communications and for diffusion of virulent soluble factors during disease onset and progression ^[2]^. Indeed, plaque topography suggests that nutrient, moisture, and oxygen gradients promote the formation of microenvironments in which the oral microbiome actively maintains healthy homeostasis ^[16]^. Thus, there is a growing need to develop experimental tool able to recreate physical and metabolic oral conditions, including oxygen and pH gradients, to accurately study host-pathogen interactions *in vitro*.

Herein, we recapitulated several *in vivo* oral features, including native architecture, oxygen gradients, and multicellular human population, in an effort to provide a cost effective, robust and reliable experimental tool for large scale longitudinal perturbation studies ^[17]^. Based on structural biopolymers as building blocks, we have developed a humanized three-dimensional gingival tissue model that recreates the native periodontal pocket and mimics the physiological oxygen tension, thus, able to support human microbiome persistence with a physiologically relevant level of microbial diversity and native spatial organization. The modulation of inflammatory markers in the presence of oral healthy microbiome suggested the humanized functional response of this model. We are anticipating these efforts will open the door to future studies to elucidate the initial interactions and balances between these two communities that are responsible for the oral tissue homeostasis or dysbiosis, that ultimately dictates the healthy or diseased tissue states.

## 2 Results and Discussion

### 2.1 A physiologically relevant tissue model

Given the importance of a three-dimensional structure in the gingiva architecture (*i*.*e*., epithelium and connective tissue interplay, mechanotransduction, complex metabolism) and the lack of appropriate experimental tools, we aimed to develop a model that is physiologically relevant and facilitates the investigation of the large-scale perturbations shown in oral disease, while being cost-effective, robust, and reliable. To successfully mimic the architecture of the gingiva, we developed an *in vitro* humanized gingiva that employs silk proteins as a sponge scaffold to support the growth of gingival and microbial cells **(Figure 1, A-B)**. Silk fibroin has been used as sutures for decades and has been recently cleared for FDA approval (SilkVoice, Sofregen Inc.). Unlike other natural polymers, silks are unique high molecular weight, amphiphilic proteins that possess remarkable strength and toughness exceeding other commonly used degradable polymeric biomaterials ^[18]^. Silk regenerated from aqueous solution can form hydrogels, fibers, sponges, films, etc., with properties tailored to mimic tissue functions ^[19]^. Silk fibroin has also shown excellent biocompatibility both *in vitro* as well as *in vivo*. The gingival scaffold was fabricated by casting a replica mold of an adult lower gingiva (**Supplementary Figure 1, A-B**) to recreate a tooth-gum unit with a porous structure to facilitate homogeneous oxygen diffusion and nutrient distribution, as shown in **Supplementary Figure 1, D**. Lastly, teeth were 3D printed with dental resin, shown to be compatible with bacterial viability and growth ^[20]^. The architecture of the scaffold **(Figure 1,C)** was optimized to mirror the healthy depth (0.69 mm) of the human dentogingival junction **(Supplementary Figure 1, C)**^[21]^, a space that lies between the gingiva and the tooth and whose increase in depth (≥4 mm) is one of the indices for assessing gum disease ^[22]^. To the best of our knowledge, this is the first gingival *in vitro* model that recreates with high-fidelity the anatomical niche where the oral host and microbiome co-exist. Previous studies have explored microbiome persistence on the apical surface of a Transwell system ^[23]^, on hydroxyapatite discs ^[15]^ or inserted into microfluidic devices ^[24]^ to establish contact with the underlying keratinocyte layer. Albeit these models were able to replicate some of the oral features (*i*.*e*., oral mucosa, cytokines secretion), they lacked the native pocket, and, more importantly, they neglected the maintenance of a normoxic to hypoxic gradient, which influences the eubiotic organization of polymicrobial communities and thereby their crosstalk with the host. Moreover, clinical studies have correlated dysbiosis with increased depth (greater than 4 mm), hypoxic (O2: 1-3%) and acidic pockets in periodontal disease ^[25] [22] [26]^. In our model, the anatomical geometry of the sulcus creates a normoxic milieu at the epithelium-aerobic population interface, while providing a hypoxic niche for facultative and strictly anaerobic bacteria towards the core of the scaffold. To prove the presence of an oxygen gradient, we manufactured two silk cellular sponges with different geometries, planar and anatomical, and populated with human primary oral cells (*see section 2*.*2*); we then compared the physical characteriation with the corresponding acellular constructs (planar and anatomical controls) **(Figure 1, D-E)**. The planar construct was designed to have a similar geometry to that of a Transwell culture system and a stable oxygen profile along the x-axis, as shown (**Figure 1, D**). Measurements in cellular anatomical sponges revealed an oxygen gradient with values comparable to the physiological range ^[27]^, with 18±3 mmHg in the outer portion of the pocket and 11±2 mmHg in the inner part **(Figure 1, E)**. Previous planar models, in fact, have recorded a loss of biofilm viability due to the lack of hypoxia gradients in their systems ^[23]^, while others have grown bacterial lines in separated aerobic and anerobic conditions, subsequently combined to support their survival ^[15]^. Our results demonstrated the formation of localized oxygen tension, as a result of the native geometry and structural properties of the gingival scaffold. Taken together, those features will provide a suitable environment for subgingival microbial communities, facilitating their spatial organization and, thereby, promoting their abundance and diversity.

**Figure 1:**
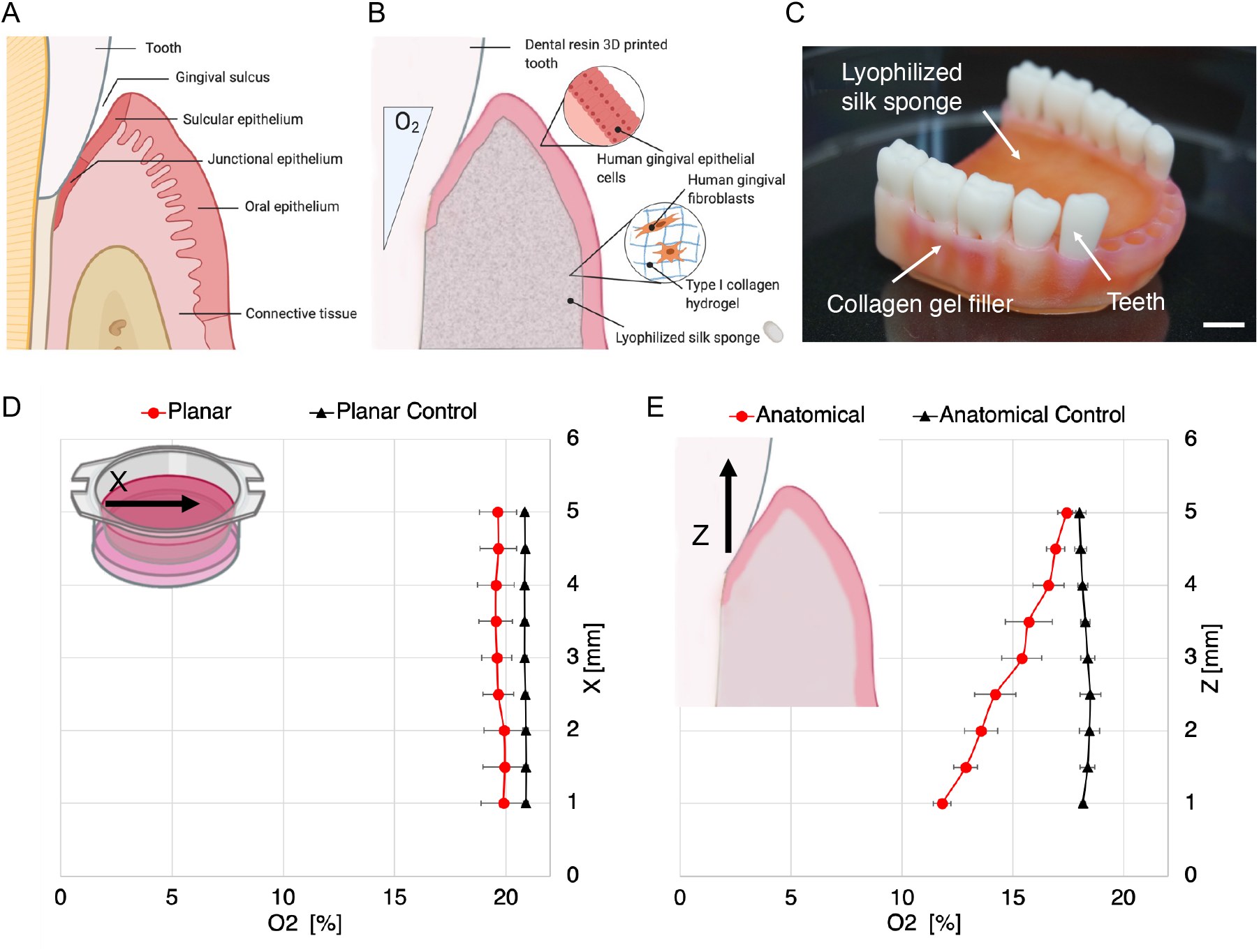
Anatomical Gingival Tissue Model. A. Schematic of the human gingival tissue with focus on the periodontal pocket. B. Schematic of the human anatomical gingival tissue model comprised of a dental resin 3D printed tooth interfaced with silk-collagen scaffold seeded with human primary gingival epithelial cells and fibroblasts. C. Macro image of the in vitro anatomical tissue model. Scale bar = 1 mm. D. Spatial oxygen profile of the cellularized anatomical model (red line) in comparison to control plain scaffold (black line) and planar tissue architecture (red line with cells, black line without cells).

### 2.2 The oral epithelium: an antimicrobial barrier

Histologically, the gingiva is composed of oral mucosa, a barrier that separates the host from its environment and protects it against infections, but also connective tissue, that holds firmly the gum to the dentition ^[28]^. Considering the physiological importance of both tissues and their crosstalk, we aimed to replicate the gingival cytoarchitecture within the scaffold **(Figure 1, B)**. To mimic the oral mucosa and submucosa, we employed human primary gingival (keratinocyte and stomal) cells; specifically, stromal cells were embedded in type I collagen extracellular matrix to provide structural integrity inside the scaffold **(Figure 1, B-C)**, while keratinocytes were seeded on the closed porosity of the scaffold. Previous gingival studies have used immortalized cell lines that can be expanded for prolonged passages and recapitulate the histological features of the gum ^[23] [15]^; however, immortalized cells have been shown to fail to mimic cellular responses to *P. Gingivalis*, a keystone pathogen in periodontal disease ^[29]^, suggesting their limited usability. To assess cellular viability in the construct, we labeled gingival cells with Calcein-AM indicators showing prolonged viability in the construct after six weeks in culture **(Figure 2, A-B)**. These findings agreed with the ability of the scaffold to support the long-term growth of keratinocyte and stromal cells. The homogeneous distribution of the pores in the scaffold, in fact, can maintain the construct viable for extended periods favoring cellular growth, differentiation, and native tissue functions, as previously shown in other *in vitro* studies conducted on intestinal ^[30]^, brain ^[31]^ or kidney ^[32]^ tissues. Subsequently, to assess native histological features, we performed scanning electron microscopy (SEM) and histology imaging analyses **(Figure 2, C-D-E)**. SEM micrographs indicated the formation of a basement membrane characterized by stromal cells, which promoted the development of a multi-layered epithelial structure lying above **(Figure 2, C-D)**. Analysis of the upper sulcular epithelium showed, in fact, the protrusion of lamellipodia in some of keratinocyte cells **(Figure 2, C*i*)** ^[33] [34]^, while the rearrangement of connective tissue with new synthesis of collagen fibrils reorganized into individual and mesh-like fibrils is shown in **Figure 2, D**. ^[35]^. Histological characterization of the 3D model revealed the formation of the sulcular portion of the oral mucosa, tightly bounded by dense underlying connective tissue **(Figure 2, E)**, indicating an organization similar to the one in the native gum ^[36] [37]^. Lastly, the epithelium displayed a physiological organization, as shown by the positive staining E-Cadherin **(Figure 3)**, a protein essential in the formation of adherent junctions between cells, to suggest the presence of epithelial barrier integrity ^[36]^. Additionally, to highlight multi-layering and differentiation within the epithelium, we counter-stained keratinocytes with Ki67, a protein that is not expressed in quiescent cells (G0 phase) ^[38]^ and thus distinguishes cells in a state of active proliferation from differentiated (quiescent) cells forming the shedding epithelium ^[23]^ **(Figure 3, B)**. Immunohistochemistry results indicated Ki67 positivity in the lower layer, while strong expression of E-Cadherin in both the lower and upper layers, supporting the epithelium barrier formation ^[39] [23]^ **(Figure 3, A)**. To further characterize the barrier integrity prior to microbiome inoculation, we quantified the Trans Electrical Epithelium Resistance (TEER) in a planar construct populated with gingival cells by means of a voltohmeter **(Figure 3, C)**. Alongside the three-dimensional construct, we used a two-dimensional culture of keratinocyte and stromal cells to corroborate the immunohistochemical data, confirming the ability of keratinocytes to form functional multilayers with enhanced barrier activity in the 3D construct. Compared with keratynocytes alone, whose TEER remained constant throughout the three weeks, the 3D construct showed an increase in TEER overtime and higher values than the 2D models, confirming the establishment of a functional barrier in the tissue model **(Figure 3, C)**. Indeed, the construct displayed after seven days physiological values of gingival epithelial multilayers ^[40]^ that were maintained stable for the entire duration of the culture. Taken together, these data confirmed the adhesion, spreading and integration of the primary gingival cells with the silk scaffold ^[41]^ and the recapitulation of the oral mucosa and submucosa *in vivo* features; moreover, they endorsed the use of silk as material for the development of long-term 3D gingival tissue.

**Figure 2:**
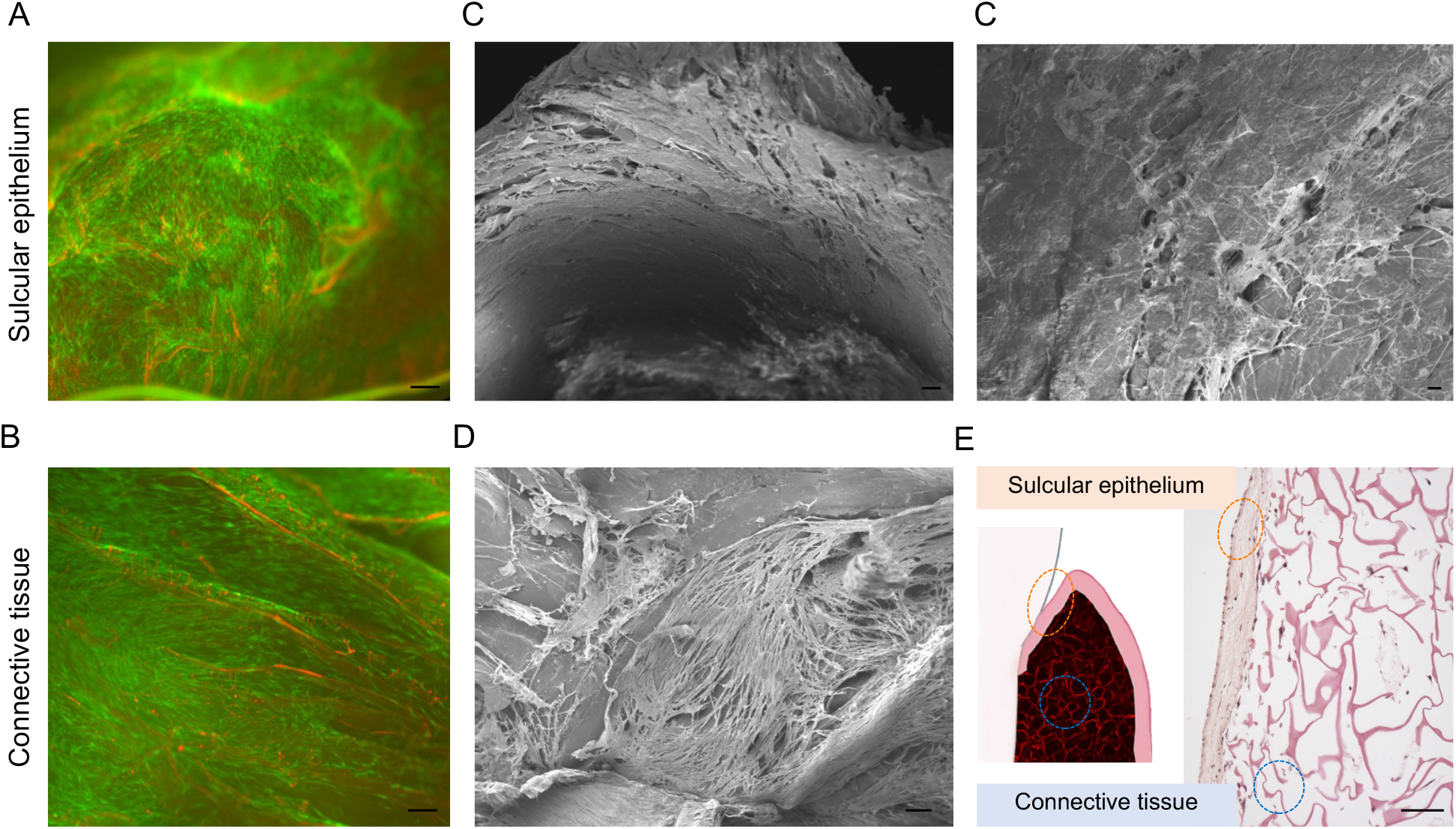
Anatomical Gingival Tissue Model Biological Assessments. Sulcular epithelium and connective tissue viability and morphological characterization at 6 weeks in culture. Maximum intensity projection of confocal laser scanning microscopy analysis of Calcein-AM labeled hGECs and hGFCs (A and B) and SEM micrographs (C and D). E. H&E staining of gingival anatomical model histological section, showing stratified sulcular epithelium and populated connective tissue. Scale bars = 100 µm.

**Figure 3:**
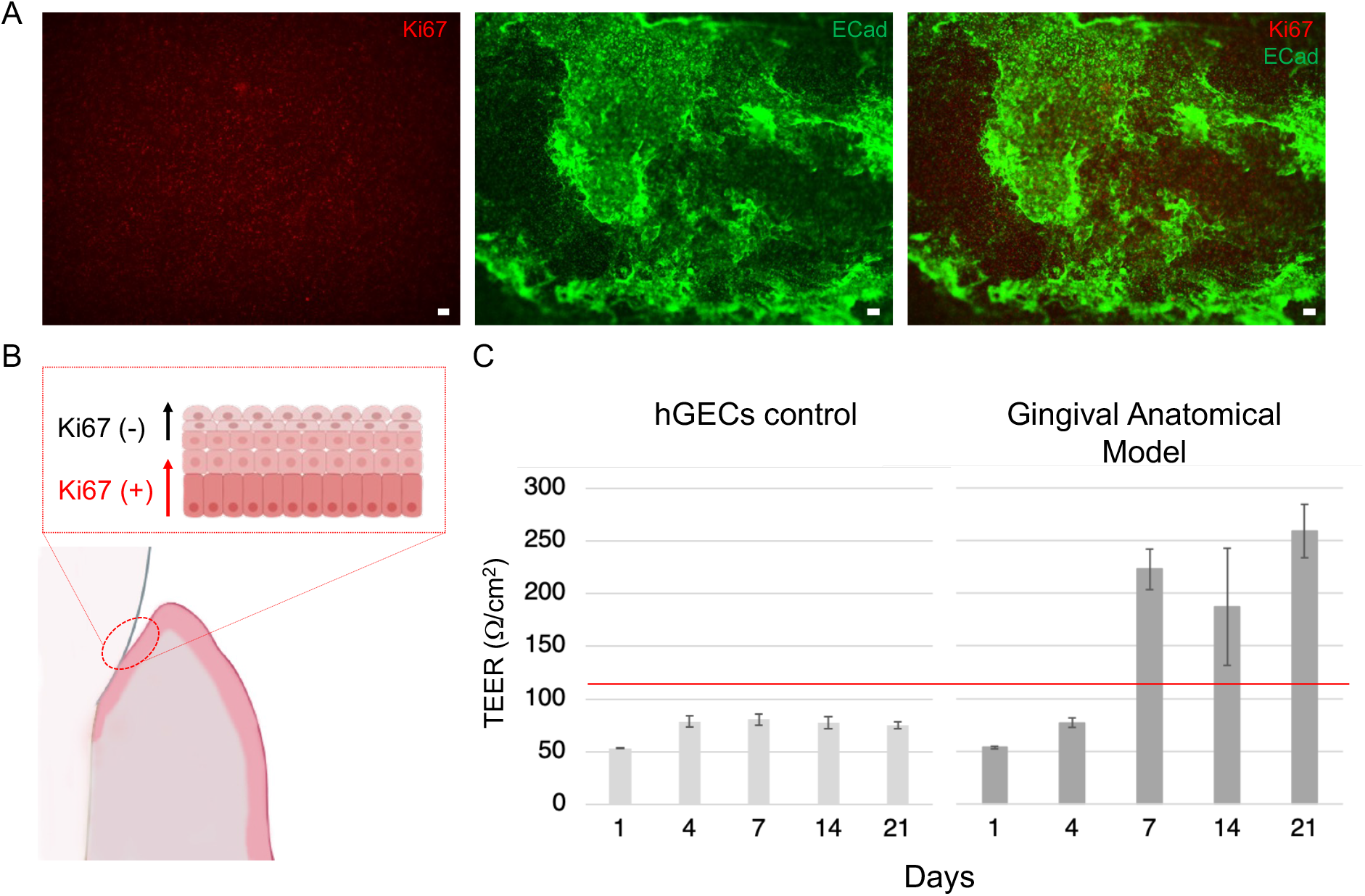
Epithelial Barrier Functional Assessments. A. IHC assessment of epithelium proliferative profile and differentiation. Maximum intensity projection of confocal laser scanning microscopy analysis of Ki67 and ECad (Scale bars = 100 µm) at 6 weeks in culture. B. Schematic of the epithelial barrier model. C. Epithelial transmembrane resistance measured over time for anatomical gingival tissue model in comparison to hGECs alone and physiological gingival tissue reference (red line) ^[76]^.

### 2.3 Maintenance of a complex human oral microbiome in vitro: viability and spatial distribution

To replicate the physiological polymicrobial dynamics within the biofilm and initiate interactions with the host, we inoculated human subgingival plaque isolated from healthy patients into the periodontal pocket **(Figure 4)**. Physiologically, the host provides a suitable ecosystem in which microbial communities can adhere, live, instruct the host immune response, and suppress the initial growth of pathogenic species to maintain homeostasis ^[2] [42]^. To support tissue homeostasis *in vitro*, we supplemented coculture media with pooled human saliva, with the aim of complementing community nutrition and pH stabilization (*i*.*e*., urea, citrate, uric acid) ^[43]^, but also to protect mammalian cells, as human saliva contains antimicrobial factors ^[42] [2]^(*i*.*e*., lysozyme or lactoferrin). Live staining with Syto9 confirmed the integration of the microbial communities and their viability after 24 hours in the construct **(Figure 4, A)**. To further characterize the microbiome in the anatomical model, we tested the hypothesis of a spatial distribution of the biofilm as a result of the oxygen content, by imaging three regions of the construct that were previously identified through oxygen measurements. SEM micrographs **(Figure 4, B)** showed a microbial spatial organization, as function of oxygen level, and aggregation into round-shaped and corncob structures in the regions of the periodontal pocket, as an indication of biofilm formation. Data in literature, in fact, indicated that the biogeography organization of the microbial cells dictates their ecology and physiology, but also their interplay with the host ^[44]^. Additionally, corncob structures have been previously identified in subgingival plaque samples as an index of physical direct contact of different genera of bacteria for assemblage or attachment (i.e., *Streptococcus* and *Aggregatibacter*) ^[44]^. Therefore, we have proved that the formed oxygen gradient in the scaffold dictated the biogeography of microbial cells.

**Figure 4:**
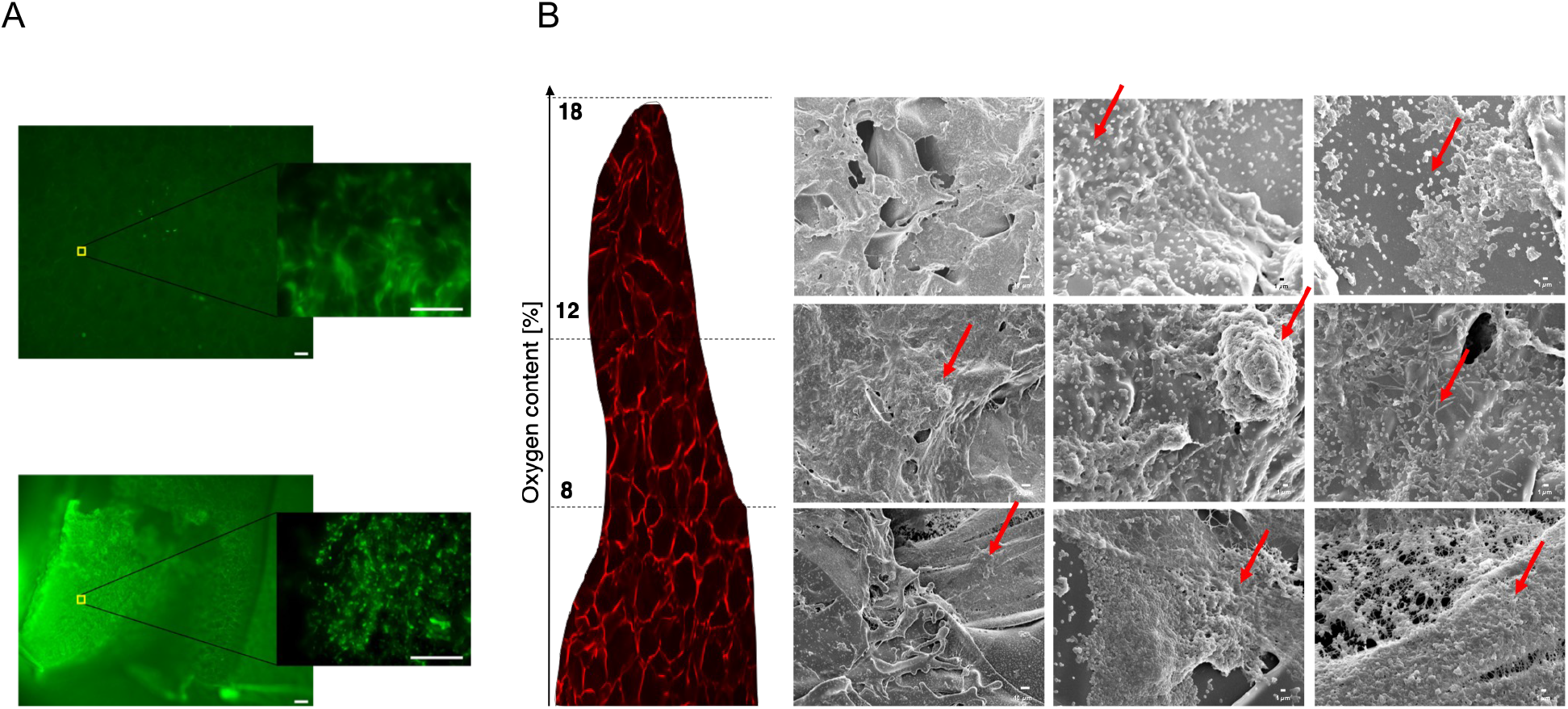
Human Oral Microbiome Viability and Organization. A. Microbiome viability assessment at 24 h in the anatomical gingival tissue model in comparison to the planar model control. Maximum intensity projection of confocal laser scanning microscopy analysis of Syto 9 positively stained human oral microbiome samples. Scale bars = 100 µm. B. Scanning electron microscopy analysis showed a gradient of plaque organization as a function of the oxygen content.

### 2.4 Microbial identity in the 3D gingival tissue model: 16S rRNA

Multiple species of bacteria inhabit the oral cavity and molecular studies have been conducted to identify the molecular bacterial signature and understand which genera are under or over-represented in eubiotic or dysbiotic scenarios. Our model aims to mimic a condition of periodontal health. To analyze the capability of the anatomical model to sustain the growth of a healthy oral microbiota (anerobic and anaerobic phyla), we performed a 16S rRNA sequencing analysis comparing the bacterial profile obtained after a one-day incubation period (24h Microbiome Anatomical Model) with the original sample (Initial Microbiome) **(Figure 5)**. Principal component analysis (PCA) **(Figure 5, A)** showed a dissimilarity in the oral microbiota between the two conditions, confirmed by the relative abundance of the phyla of the microbial populations **(Figure 5, B)**. Compared with Initial Microbiome, we found that the anatomical model supported the viability and growth of the complex human microbiota. Indeed, of the nine Operational Taxonomic Units (OTUs) identified in the Initial Microbiome as phyla, eight were retained in the anatomical model. In particular, we found that phylum *Proteobacteria*, which is an health associated phyla in periodontal health ^[45]^, was highly enriched after one day of incubation in the anatomical model **(Figure 5, B)**, while some phyla were reduced, especially *Actinobacteria, Saccharybacteria* and *Synergistetes*, or even disappeared such as *Spirochaetes* (**Figure 5, B)**. Although we could not mimic the exact relative abundance found in Initial Microbiome one day after inoculation, we were able to preserve the majority of the richness and diversity with aerobic and anaerobic species coexisting in the anatomical model. In fact, we were able to identify a total of 294 OTUs, including 99 OTUs shared among the samples **(Figure 5, C)**. Additionally, both the α-diversity, as determined by Shannon index, and the richness, as determined by Chao-1 index, showed significant statistical reduction from the original sample in the anatomical model **(Figure 5, D-E)**. Both reductions could be associated to short-term culture period of the model. In fact, i*n vitro* studies conducted on the gut microbiome have reported an initial reduction in richness and diversity that stabilizes after a prolonged period in culture ^[46] [47]^. Furthermore, to better comprehend which OTUs were retained in the anatomical model, we characterized the microbiome as a genus in the taxonomic rank **(Supplementary Figure 2)**. The analysis revealed a total of 82 genera, among which the well-characterize *Streptococcus, Actinomyces, Fusobacterium, Neisseria, Porphyromonas, Prevotella, Capnocytophaga, Rothia, Leptotrichia and Veillonella* ^[48] [49] [50]^. Compared against Initial Microbiome, we observed decrease relative abundance in the anatomical samples for *Actinomyces, Porphyromonas, Prevotella, Rothia, Leptotrichia and Veillonella*, while similar levels of relative abundance for *Fusobacterium*, and increase relative abundance of *Streptococcus, Neisseria* and *Capnocytophaga* in the anatomical samples. Interestingly, *Streptococcus* and *Fusobacterium* are among the major bacterial genera which colonize the oral cavity and form crucial constituents of dental plaque (i.e., dental biofilms accumulating on non-shedding tooth surfaces). *Fusobacterium* members, in fact, act as a bridge between early (*Streptococcus*) and late colonizers, coaggregating with most oral bacteria ^[51] [52]^. These findings correlate with the *in vivo* tissue-specific tropisms mimicked in the anatomical model, in which the presence of colonizers facilitates structural polymicrobial synergies and thus colonization and aggregation of other bacteria on oral surfaces ^[2]^. Lastly, to further analyze changes in the relative abundance of the microbiota between the Initial Microbiome and the anatomical model, we, generated a volcano plot and characterized the microbiome as a species in the taxonomic rank. The volcano plot represents the fold change in the relative abundance; specifically, a negative fold change less than -2 denotes species enrichment in the hPlaque0 samples (white circles), whereas a positive fold change greater than 2 denotes species enrichment in the anatomical model samples (black circles) with a 95% confidence interval **(Figure 6, A)**. A total of 50 species showed a statistical difference in relative abundance, 42 corresponding to the Initial Microbiome samples and 8 corresponding to the anatomical model **(Supplementary Table 1)**. The 8 species with the highest abundance were identified as members of the genus *Neisseria* and *Gemella*, a member of the order *Lactobacillales*, and the species *Neisseria subflava, Streptococcus constellatus, Solobacterium moorei, Fusobacterium nucleatum subsp. polymorphum and Capnocytophaga sputigena*. Interestingly, of these species, only *N. subflava* is considered aerobic, while the others are considered either facultative anaerobes or strict anaerobes, like *F. nucleatum subsp. polymorphum*. In comparison to the Initial Microbiome, we observed an increase in the relative abundance of all species in the anatomical samples, which in the case of *F. nucleatum subsp. polymorphum* and *Capnocytophaga sputigena* were statistically significant **(Figure 6, B-F)**. With regard to the statical difference of some species between the two conditions, this could be associated with several factors, including, but not limited to, saliva supplementation, as a direct effect (i.e., environmental factors, personal hygiene, sex or periopathogens) ^[53] [54] [55]^ on the composition of the subgingival microbiome, but also experimental conditions, such as components of co-culture media or teeth for attachment, that could cause some species to be more selective than others. Altogether, these results demonstrated that the proposed 3D anatomical gingival model can support the survival and growth of key members of the healthy human oral microbiota (aerobic, facultative and strictly anaerobic), including uncultivable bacteria (i.e., *Saccharibacteria*-TM7) ^[56] [57] [22]^.

**Figure 5:**
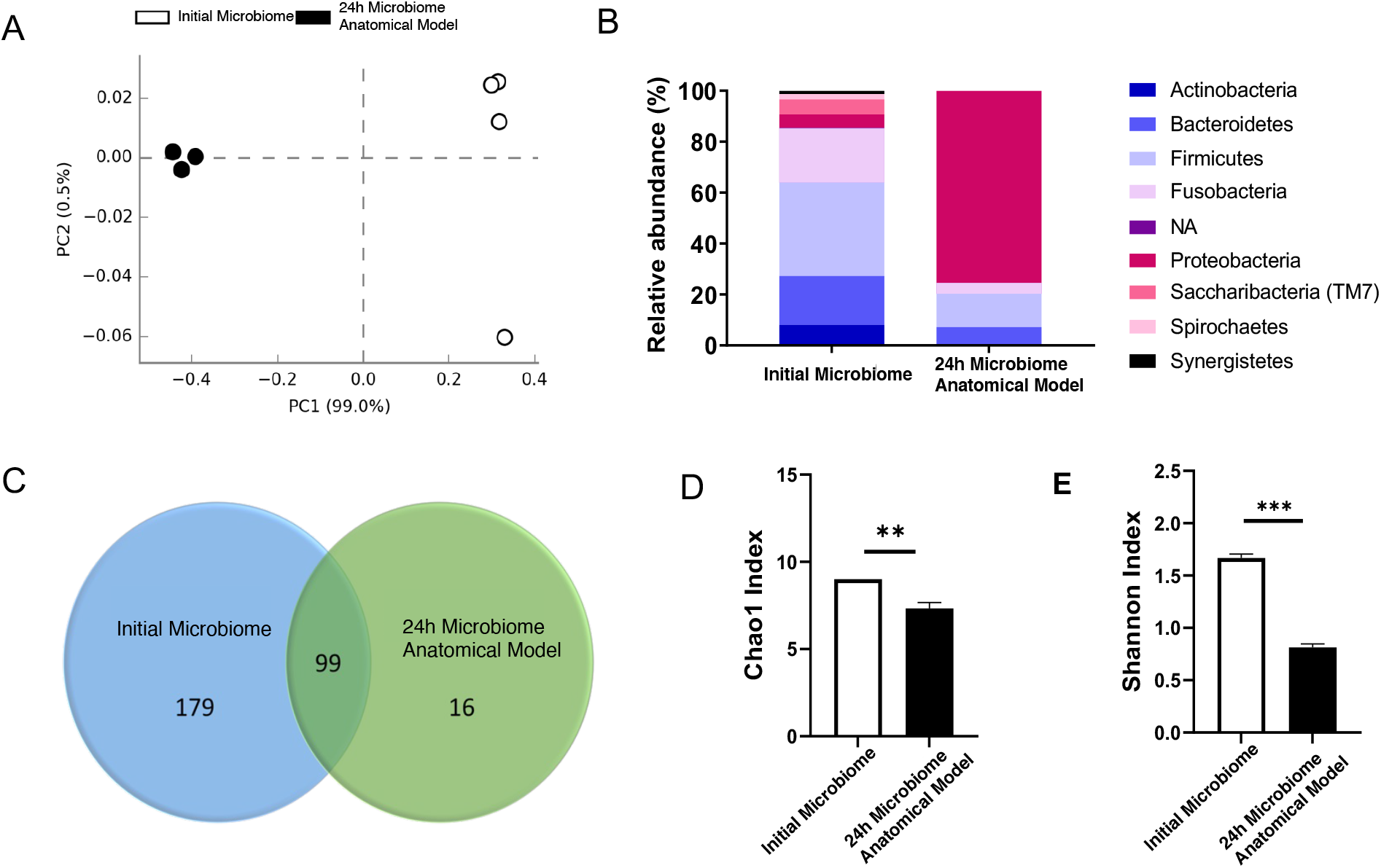
Analysis of the diversity and relative abundance of human oral microbiome at 24 h. A) Principal component analysis (PCA) clustering of oral microbial populations generated after 16S rRNA sequencing of samples collected from human plaque (hPlaque0) and after incubation in the anatomical model for 1d (Anatomical). Each point represents one sample. B) Relative abundance distribution of main phyla detected in the human plaque before incubation (hPlaque0) samples and samples collected after 1 day incubation in the anatomical model (Anatomical). C) Venn diagram showing unique and shared OTUs D) Alpha diversity representation of oral microbiota. F) Richness representation of oral microbiota. Statistical analysis: ANOVA test with a t-Student posthoc test. *p < 0.05; **p < 0.01; ***p < 0.001.

**Figure 6:**
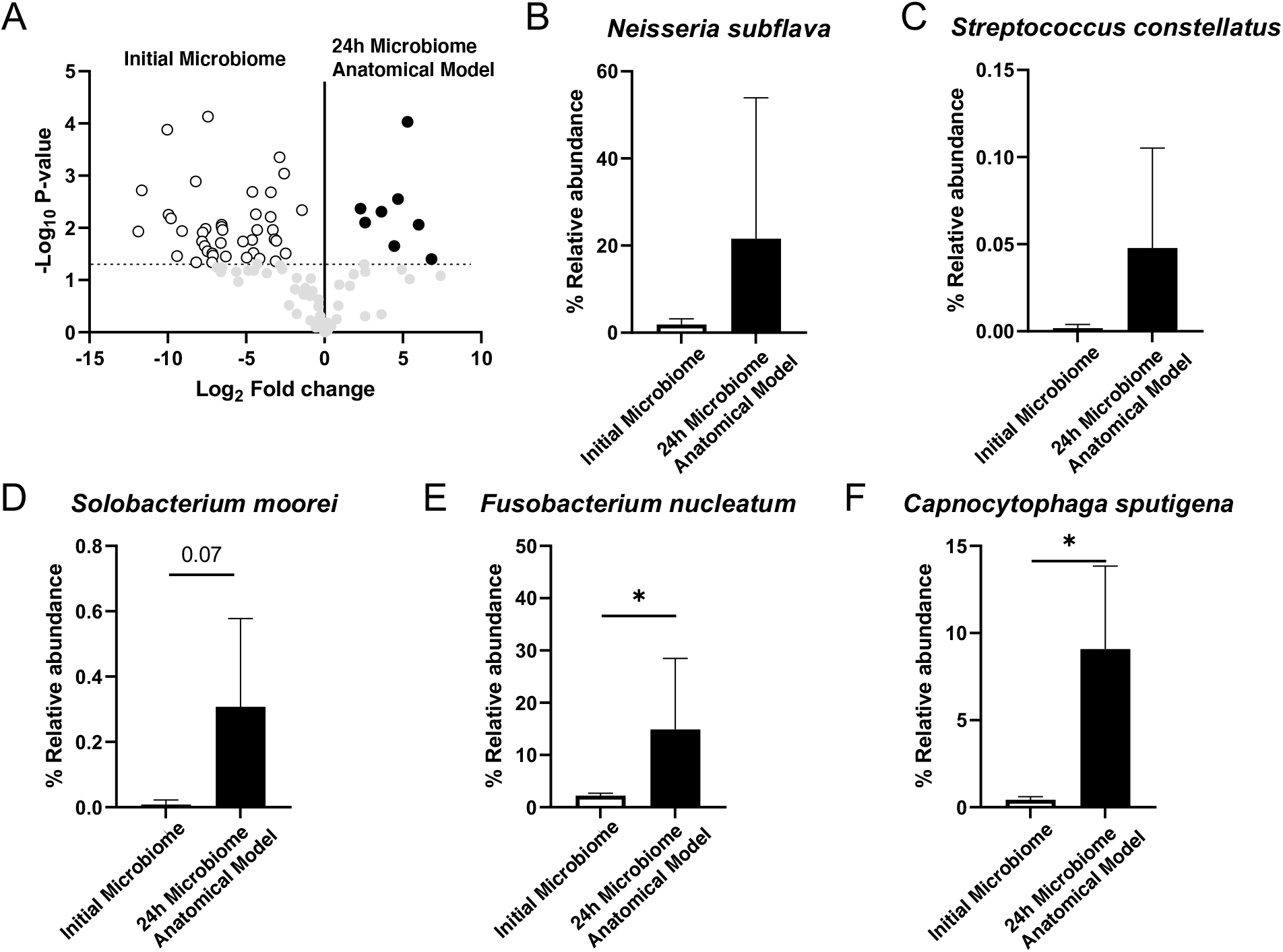
Ad-hoc Specie Analysis of human oral microbiome at 24 h. A) Volcano plot of specie level. The x axes display the fold changes in relative abundance (log2) between original oral plaque samples (hPlaque0) and after 1d incubation in the anatomical model (Anatomical) data sets; the y axes display the log of the P values of the test statistic. The dashed horizontal line at log 1.3 corresponds to a P value of 0.05. Enrichment of the specie in the hPlaque0 samples is represented as white circles, whereas enrichment of the specie in the anatomical model samples is represented with black circles. The full list of species showing significant shifts in relative abundance is provided in Supplementary Table 1 in the supplementary section. B-F) Relative abundance of Neisseria subflava (B), Streptococcus constellatus (C), Solobacterium moorei (D), Fusobacterium nucleatum subsp. polymorphum (E) and Capnocythophaga sputigena (F). Statistical analysis: ANOVA test with a t-Student posthoc test. *p < 0.05.

### 2.5 Tissue response to human oral microbiome: cytokine profile

Considering the proximity of commensal bacteria to the oral mucosa, the host implements surveillance mechanisms to maintain tissue homeostasis and train the immune systems, such as infiltrating neutrophils or resident immune cells ^[58] [59]^. It may be argued that, under healthy conditions, the immune response induced by the oral microbiome is appropriate and the release of pro/anti-inflammatory cytokines is moderate ^[60]^; on the other hand, the initiation of a dysbiotic state by pathobionts and the consequent tissue disruption leads to over-activation of the immune system and, therefore, a positive-feedback loop of pro-inflammatory cytokines ^[59]^. To validate the clinical relevance of the 3D gingival model, we investigated the initial response of the anatomical tissue model to the human microbiome inoculated from healthy patients. In both samples, pro-inflammatory (GM-CSF, IL-1RA, IL-1*α*, IL-1β, IL-6, IL-8, IL-12p40, IL-17A, and TNF-*α*) and anti-inflammatory (IFN-*γ*, IL-2, IL-10, IL-3, and IL-4) cytokines were above the detection limits, except for IL-10, IL-1RA, IL-3 and IL-4 **(Figure 7)**. In comparison to the untreated anatomical tissue model, the inoculated anatomical group showed a decrease of both pro-inflammatory and anti-inflammatory cytokines **(Figure 7)**. Cytokines downregulation has been previously reported as part of the initial response of the gingival epithelium to interactions with oral microbiome ^[61] [60]^. For example, IL-1β and IL-8 are statistically reduced in presence of the microbiome in the anatomical model compared to the untreated **(Figure 7)**, as shown in *in vivo* findings, where moderate plaque formation is considered in the normal range of surveillance mechanisms by the host ^[60]^, while increased cytokines release is associated with inflammatory gingival conditions ^[61] [62]^. Overall, the model supported a functional response to human oral microbiome interactions in healthy conditions.

**Figure 7:**
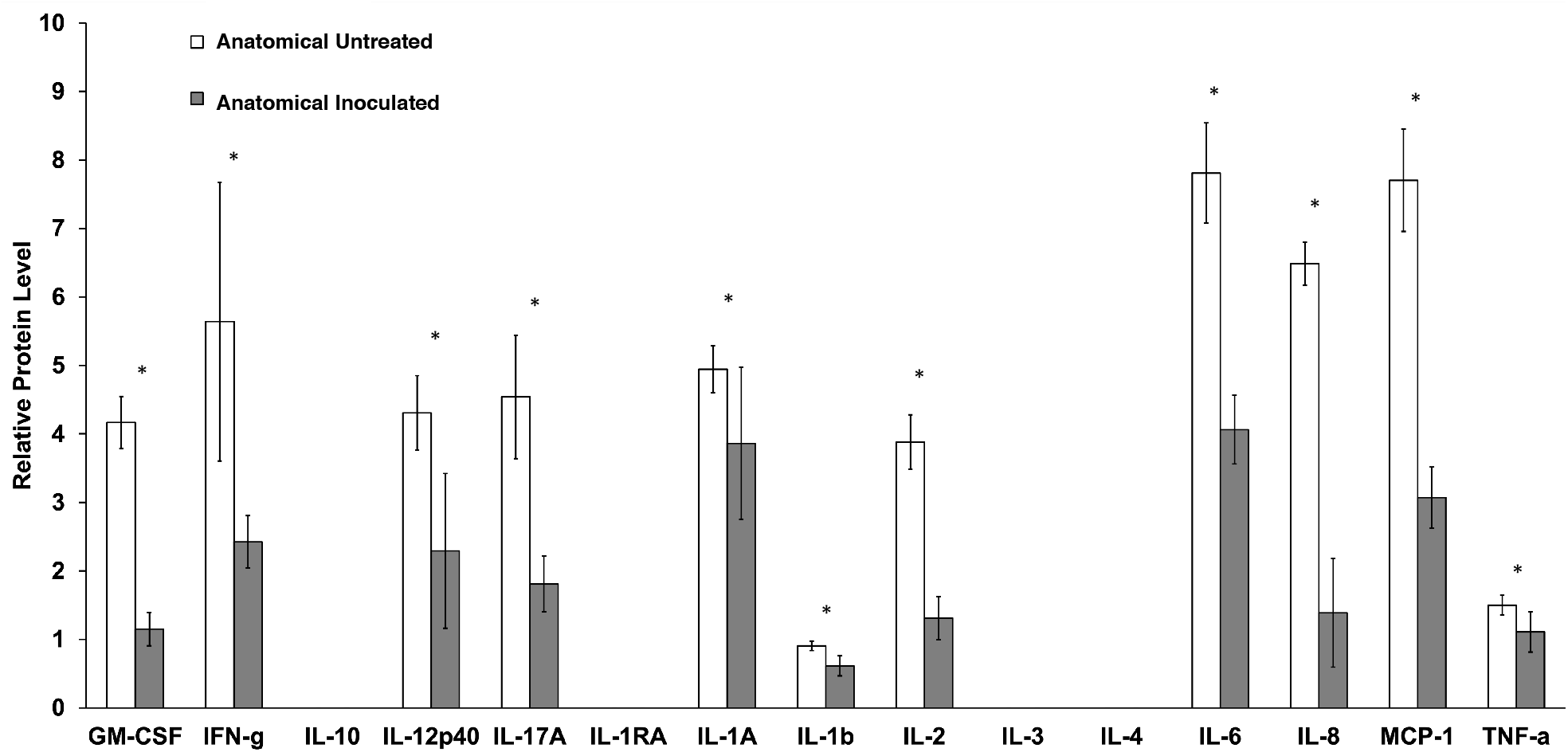
Tissue response to human oral microbiome: cytokine profile. Simultaneous analysis of multiple cytokine and chemokine biomarkers with Bead-Based Multiplex Assays using the Luminex technology at 24 hours upon inoculation in cell culture samples. * significant effect of human microbiome samples on the initial inflammatory response of the 3D gingival anatomical tissue model (p<0.05).

## 3 Conclusion

The aim of this study was to develop an *in vitro* humanized gingival model that mimics the *in vivo* anatomical and cytoarchitectural features of the gingiva and host-microbiome interactions. It is hypothesized that chronic periodontitis results from gingivitis progression ^[3]^, current technologies are limited and failed to sustain and recapitulate the complex host-pathogen interplay under healthy and disease conditions. To the best of our knowledge, no other oral tissue model has been able to mimic the gingival anatomical architecture and the physiological oxygen tension that, together with oral mucosa stratification, allow aggregation and organization of the microbiome while maintaining diversity *in vitro*. Overall, our results indicated the establishment of a tissue phenotype, maintenance of an eubiotic microbiome after one day of culture, and inflammatory signaling similar to that *in vivo*. In conclusion, these findings proved the applicability of the gingival model as a platform for clinical investigations of periodontal health and disease.

## 4 Experimental Methods

### 4.1 Tissue model fabrication

The scaffolding process used a water-based silk technology to create highly porous silk scaffolds for implants and tissue regeneration ^[63]^. Highly interconnected pore architectures were achieved in the silk scaffolds by freezing and lyophilizing the water in the silk polymer solution.

#### 4.1.1 Silk solution preparation

Aqueous silk solution was prepared from *Bombyx mori* silkworm cocoons, following the experimental procedure described in previous studies ^[64]^. *B. mori* silk cocoons were purchased from Tajima Shoji Co. (Yokohama, Japan). Briefly, the cocoons were degummed by boiling in 0.02-M sodium carbonate (Sigma Aldrich, St Louis, MO) solution for 30 min. The extracted fibroin was then rinsed three times in Milli-Q water, dissolved in a 9.3-M LiBr solution yielding a 20% (w/v) solution, and subsequently dialyzed (MWCO 3,500) against distilled water for 2 days to obtain silk fibroin aqueous solution at the approximate concentration of 8% (w/v), as determined by gravimetrical analysis.

#### 4.1.2 Polydimethylsiloxane (PDMS) molds preparation

The silk anatomical gingival scaffolds were prepared using a replica molding technique **(Supplementary Figure 1)**. The design of a human adult gingival anatomical model and relative teeth were purchased from CadHuman.com and used to 3D print the lower gum structure with a Form2 printer in combination with a white dental resin (formlabs, Somerville, MA, USA). Sylgard 184 silicone elastomer kit was prepared in a 10:1 ratio, where the 3D printed model was inserted, and placed for 3 hours in a 70°C oven to cure.

#### 4.1.3 Silk scaffold preparation

Upon model retrieval, the resulted anatomical mold was used to cast aqueous silk solution (% wt/v in deionized water), while planar control scaffolds were prepared by dispensing into wells of standard 24-well cell culture plates (1 mL/well) (Corning, NY). Dry scaffolds were removed from the molds and rendered insoluble in aqueous environments by autoclaving at 121°C for 20 min at 15 psi to induced β-sheets formation. Scaffolds were rehydrated in deionized water. Planar scaffolds were sliced with a micron cutter using a razor blade. Anatomical and planar scaffolds were then sterilized by autoclave in deionized water before cell culture procedures.

### 4.2 Anatomical and planar tissue models preparation

#### 4.2.1 Cell culture procedures

Human gingival epithelial cells (hGECs) and human gingival fibroblast cells (hGFCs) were purchased from Lifeline Cell Technology (Frederick, MD), derived from the lower jaw gingiva. Gingival cells were maintained in culture up to passage 6 in appropriate medium supplemented with associated growth factor kits (Lifeline Cell Technology, Frederick, MD). For passaging cells, both cell types were detached using 0.05% Trypsin/0.02% EDTA coupled with Trypsin Neutralizing Solution (Lifeline Cell Technology, Frederick, MD) and centrifuged at 150 x g for 5 minutes. hGECs have been tested positive for cytokeratins K13 and K14. For the anatomical model, the culture conditions were optimized to support the growth of both hGECs and hGFCs at the same time, by using three parts of basal media for hGFCs and one part for hGECs and supplementing with growth factor kits; specifically: human serum albumin (500 µg/mL), linoleic acid (0.6 µM), lecithin (0.6 µg/ml), recombinant human fibroblasts growth factors (5 ng/ml), recombinant human epidermal growth factor (5 ng/ml), recombinant human transforming growth factor beta 1 (30 pg/ml), recombinant human insulin (5 µg/mL), ascorbic acid (50 µg/mL), L-glutamine (7.5 mM), hydrocortisone hemisuccinate (1 µg/mL), epinephrine (1 µM), apo-transferrin (5 µg/mL), recombinant human transforming growth factor alpha (0.5 ng/ml), gentamicin (30 µg/mL), and amphotericin (15 ng/ml).

#### 4.2.2 Tissue model preparation: anatomical and planar

The surface of the anatomical scaffolds was used to accommodate hGECs, while the porous bulk space housed hGFCs. hGFCs were seeded into the anatomical scaffold via neutralized rat tail collagen type I gel (First Link UK Ltd.) at 200,000 cells/ml to allow uniform distribution of the cell population. Collagen gel solution was made by mixing four parts of Collagen I and one part of Dulbecco’s Modified Eagle’s Medium (DMEM) 10X (Sigma Aldrich, St Louis, MO) and neutralized in Sodium Hydroxide 10M (Sigma Aldrich, St Louis, MO). After gelation time (37°C for 30 minutes), the construct was seeded with hGECs at a density of 50,000 cells/cm^2^. After 2 hours in incubation at 37°C, the construct was flipped and seeded on the other side with hGECs and incubated for another 2 hours. During the incubation, a small amount of co-culture medium will be dripped on the scaffolds to keep them moist. The cellularized constructs were then incubated for 1 week submerged in co-culture media until the hGECs reached confluence. Dental resin teeth were inserted in the gingival pockets after a week post-seeding and the level of media was adjusted so that hGECs close to the periodontal pocket were exposed to air liquid interface (ALI) to induce differentiation, while maintaining the base of the sponge in media. The gingival model was maintained in a humidified incubator at 37°C with 5% CO2. Culture media was changed every day for the entire duration of the experiments. Planar tissue model was made for model optimization. Planar silk scaffolds were made as described above. hGFCs in neutralized Collagen type I were seeded on the open porosity of the scaffold to recreate the bulk of the tissue. After gelation, the tissue model was flipped and accommodated inside a Transwell insert (Corning, NY) and hGECs were seeded on the closed porosity part. Culture conditions were maintained as the one for the anatomical models, dentition excluded.

### 4.3 Tissue model characterization

#### 4.3.1 Transepithelial electrical resistance

As part of the functional assessments, the gingival tissue was screen for epithelial barrier function with transepithelial electrical resistance (TEER). TEER measures nondestructively the integrity of tight junction dynamics in epithelial cell culture models, and it is a strong indicator of the integrity of the cellular barriers ^[65]^. EVOM^2^, Epithelial Volt/Ohm Meter was used with an EndOhm Chamber (World Precision Instrument) to measure TEER of the gingival construct after maturation in comparison to hGECs cultured for 7 days at ALI in a Transwell system (Corning, NY). TEER values were reported subtracting the contribution from the Transwell together silk scaffold with Collagen gel for the gingival construct and the Transwell chamber alone for the hGEC control.

#### 4.3.2 Immunofluorescence microscopy

Gingival constructs were washed with phosphate Buffer Saline-1x(PBS) (Thermo-Fisher Scientific, Waltham, MA), fixed with 4% paraformaldehyde (Electron Microscopy Sciences, 157-4) and subsequently washed with additional PBS. Permeabilization of cells was accomplished with 0.25% Triton X-100 (Sigma, T8787), followed by incubation in blocking buffer containing 1% BSA (Sigma, A4503) and 10% horse serum at room temperature (Invitrogen). Primary antibodies against Ki67 (Abcam, ab15580, dilution 1:100), E-cadherin (Abcam, ab1416, dilution 1:50) were added and incubated overnight at 4°C, followed by multiple PBS washes. Constructs were then incubated with secondary fluorescent antibodies (Life Technologies) at room temperature and washed with PBS. Microscopy was performed with a Keyence BZ-X700 microscope to image samples under 10× magnification obtained at 470–510 nm excitation over an emission range of 525–575 nm for green fluorescent protein and 560–600 nm excitation over an emission range of 630–705 nm for Texas Red (Keyence, Elmwood Park, NJ).

#### 4.3.3 Oxygen tension

The oxygen concentration profiles were measured using a PC-controlled Microx TX3 oxygen meter (PreSens Precision Sensing GmbH, Rengensburg, Germany) equipped with a needle-type housing fiber-optic oxygen sensor (NTH-PSt1-L5-TF-NS40/0.8-OIW, 140 μm fiber tapered to a 50 μm tip in the sulcus. The needle probe was mounted on a custom-made micromanipulator capable of precisely positioning the measurement spot in the vertical direction. One complete turn of the screw knob resulted in 0.1 inch (2.5 mm) of travel. Oxygen concentration was measured weekly with a 500 µm step over the scaffold profile to monitor the oxygen profile within the sulcus over 6 weeks in culture.

### 4.4 Physiologically relevant gingival tissue model: the microbiome

#### 4.4.1 Coculture with human oral microbiome in the anatomical tissue model

The ability of the anatomical gingival tissue model to maintain and support the organization of complex human oral microbiome *in vitro* was validated by inoculating pooled plaque samples (n=5) from healthy patients and cultured for 24 hours. Human subgingival plaque and saliva samples were collected from donors with healthy periodontal tissues at the Center for Clinical and Translational Research at the Forsyth Institute, Cambridge (MA). 6 plaque samples were combined and inoculated into the gingival pocket in 3 µl aliquots. Saliva samples were pooled, then sterile filtered and combined with co-culture medium at a 1:4 ratio, as previously reported ^[66] [67]^. All methods were carried out in accordance with a protocol approved by the Institutional Review Board of Forsyth (Protocol No. FIRB# 18-06), of Tufts University (Protocol No. IRB -12860) and of University Massachusetts Lowell (Protocol No. IRB – 20-090). All subjects signed FIRB approved informed consent prior to sampling.

#### 4.4.2 Morphological analysis

The morphology and distribution of construct pores, cellular morphology and organization, and microbiome spatial organization and distribution were characterized by scanning electron microscopy (SEM). The constructs were harvested, washed in PBS (1x) (Thermo-Fisher Scientific, Waltham, MA), and then fixed in 4% paraformaldehyde–0.1 m sodium cacodylate solution overnight at 4°C. After washing with deionized distilled water, samples were dehydrated at 4°C through sequential exposure to a gradient of ethanol, and then processed with a critical point dryer (Tousimis Autosamdri, USA), sputter coated with Au/Pd (Hummer VI Sputter Coater, Ladd Research Industries, USA) and analyzed by SEM (Supra55VP, Zeiss, Oberkochen, Germany) at 5 kV and 10 μA.

#### 4.4.3 Viability assessment

SYTO® 9 (Thermo-Fisher Scientific, Waltham, MA) was used to fluorescently stain nucleic acid of the human oral microbiome pooled samples to investigate viability at 24 h in the anatomical gingival tissue model in comparison to the planar construct. Before inoculation, the pooled samples were incubated for 15 min in a PBS (1x) solution of 20 µM SYTO® 9, then spun, washed, and resuspended in culture medium for seeding. Samples were then imaged at 24h post inoculation using a Confocal Laser Scanning Microscope (CLSM) with excitation at 488 nm and emission at 499–537 nm.

### 4.5 16S rRNA sequencing analysis

#### 4.5.1 Bacteria DNA extraction

Samples collected at 24 h were processed for DNA isolation using the Epicentre MasterPure Gram Positive DNA Purification Kit (Lucigen, Middleton, WI, USA). Briefly, samples were resuspended in TE buffer and incubated over night at 37°C after addition of the Ready-Lyse lysozyme solution. Then, GP Lysis solution was added, and the samples were vortexed for 1 min followed by addition of Proteinase K and incubated at 65°C for 15 minutes. Protein and RNA were then removed using MPC Protein Precipitation Reagent and RNaseA respectively. Lastly, DNA was precipitated with isopropanol (Sigma Aldrich, St. Louis, MO, USA), washed 2 times with ethanol 70% (Sigma Aldrich, St. Louis, MO, USA) and finally resuspended in water.

#### 4.5.2 16S library preparation

Bacterial 16S rRNA gene targeted amplicon sequencing was performed using a custom dual-index protocol ^[68]^. The custom 16S primers used amplified the V1-V3 region of the 16S rRNA gene and are designed to provide the best coverage of the 16S gene while maintaining high sensitivity. The sample libraries were prepared using a 22 cycle PCR reaction to reduce chimera formation. The final PCR products were purified using Ampure XP beads (Beckman Coulter, Brea, CA, USA), pooled in equal amounts, and gel purified using the QIAGEN MinElute Gel Extraction Kit (QIAGEN, Hilden, Germany). Purified, pooled libraries were quantified using the NEBNext Library Quant Kit (New England Biolabs, Ipswich, MA, USA) for Illumina.

#### 4.5.3 Sequencing and data analysis

Final libraries were sequenced on Illumina® MiSeq™ with a v2 reagent kit (Illumina, San Diego, CA, USA) (500 cycles) at the Human Oral Microbe Identification using Next Generation Sequencing core at the Forsyth Institute. The sequencing was performed at a 10pM loading concentration with >20% PhiX spike-in. For analysis, the DADA2 R package ^[69]^ was used to identify and quantify amplicon sequencing reads on the fastq files obtained after demultiplexing with the Illumina MiSeq software. Briefly, reads were trimmed and filtered to remove sequences with low quality. Quality of the trimmed and filtered reads was assessed using FastQC ^[70]^. Samples with read count smaller than 1000 reads per sample were excluded in the analysis. Results of FastQC were compiled using MultiQC ^[71]^. The trimmed and filtered reads were then processed through the denoising, concatenating read1 and read2 with a 10N spacer, and chimera removal steps of DADA2 to identify and quantify true amplicon sequence variants (ASV) present in the sample. Taxonomy of the identified ASVs was assigned using the RDP classifier algorithm ^[72]^ implemented in the DADA2 package with a training dataset developed at the Forsyth Institute and based on the eHOMD ^[73]^ generating a table with relative abundance of Operational Taxonomic Units (OTUs).

#### 4.5.4 Diversity and Richness indexes, changes in relative abundance, principal component analysis (PCA)

To calculate alpha diversity (Shannon index) and richness in the samples (Chao-1), the OTUs with specie frequency from all samples were imported into the free software Past3.2 (https://folk.uio.no/ohammer/past/). To analyze changes in relative abundance and PCA, normalized OTU tables for human plaque at time 0, planar and anatomical samples at 24h, were imported into the open-source software STAMP v2.1.3 ^[74]^.

### 4.6 Cytokine profile

#### 4.6.1 Milliplex cytokines analysis

Epithelium exudates were collected from the periodontal pocket in the inoculated anatomical models in comparison to non-inoculated samples at 24 hours. Simultaneous analysis of multiple cytokine and chemokine biomarkers for inflammation (Granulocyte-macrophage colony-stimulating factor (GM-CSF), Interferon gamma (IFN-*γ*), Interleukin 10 (IL-10), Interleukin 12p40 (IL-12p40), Interleukin 17A (IL-17A), Interleukin 1 Receptor antagonist (IL-1RA), Interleukin 1 alpha (IL-1*α*), Interleukin 1 beta (IL-1β), Interleukin 2 (IL-2), Interleukin 4 (IL-4), Interleukin 6 (IL-6), Interleukin 8 (IL-8), Monocyte Chemoattractant Protein 1 (MCP-1), Tumor Necrosis factor alpha (TNF-*α*)) was carried out with Human Cytokine/Chemokine Magnetic Bead Panel (MilliPlex MAP, Millipore, Sigma). Median fluorescence intensity of each analyte was read using a MAGPIX system (Luminex). Concentrations of proteins of interest were calculated using the median fluorescence intensity and the standard curve of each analyte, as previously described ^[75]^. Resulting protein concentrations from multiplex analyses were normalized by their respective sample DNA content.

### 4.7 Statistical analysis

For statistical analysis and dynamic identification of microbiome differences by relative abundance between the groups, STAMP v2.1.3 software was used. A Welch’s t-test for unequal variances was used to assign p-values to the OTUs when comparisons between two groups were done; ANOVA test was used when three groups were analyzed. Values of *p<0.05; **p<0.01 and ***p<0.001 were considered statistically significant.

## 5 Data availability

The main data supporting the findings of this study are available in the Article and Supplementary Information. The raw data generated in this study are available from the corresponding author on reasonable request.

## 6 Acknowledgements

Research reported in this publication was supported by the National Institute Of Dental & Craniofacial Research of the National Institutes of Health under Award Number R03DE030224. The content is solely the responsibility of the authors and does not necessarily represent the official views of the National Institutes of Health. We are also thankful to the ORAU (Ralph E. Powe Junior Faculty Enhancement Awards) for supporting this work.

## Supporting Information

**Supplementary Figure 1:**
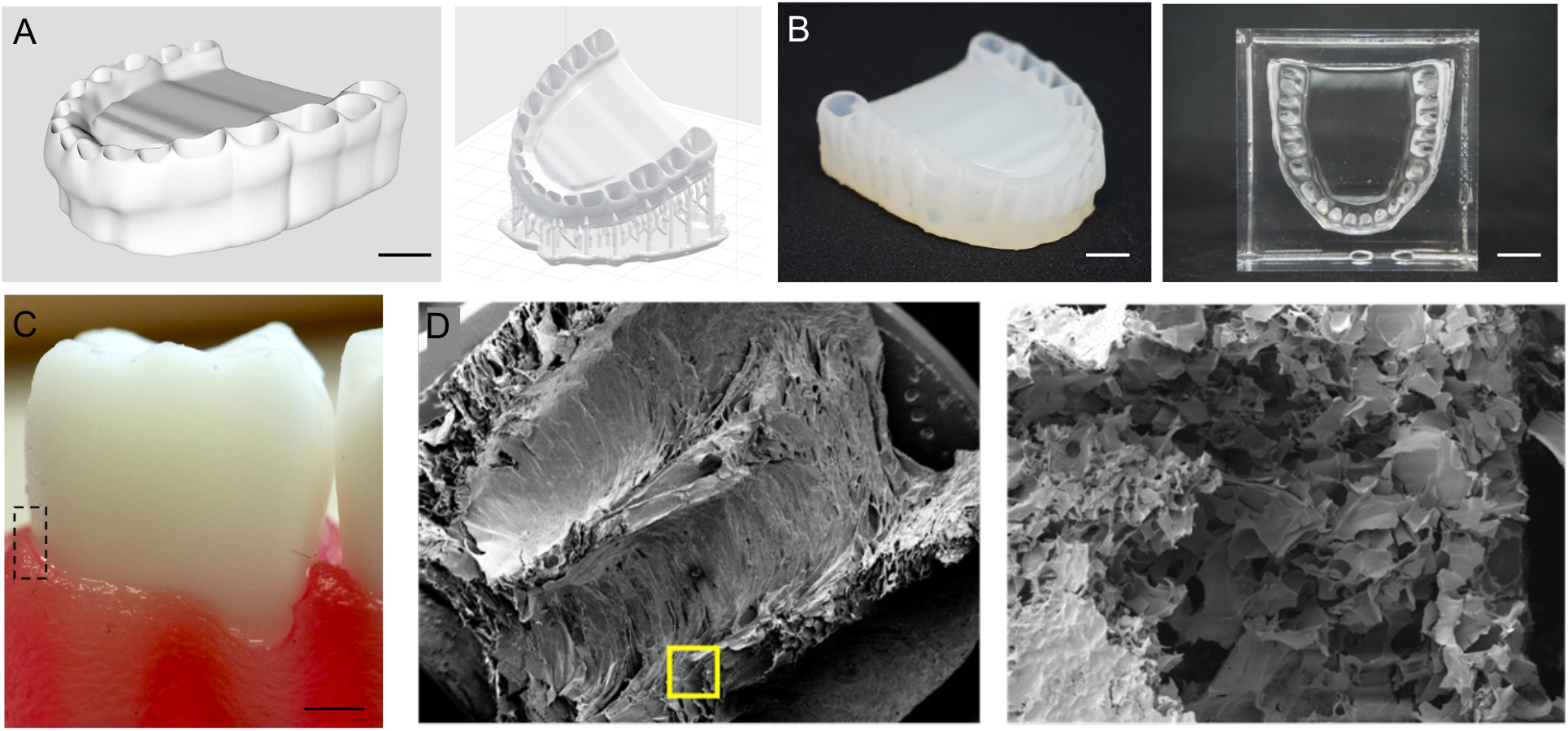
Anatomical Gingival Tissue Model Fabrication. A. Form2 3D printer specification design and rendering of the adult full mouth gingival replicate. B. 3D printed adult gingival replicate and replica mold of the 3D printed structure in PDMS. C. Macro image of the gingival pockets where the teeth are inserted during the culture to recreate the gingival pocket microenvironment. Scale bars = 3 mm. D. SEM micrographs of tissue model silk scaffold with details of the gingival sulcus (closed porosity) and connective structure (open porosity). Scale bars = 200 and 30 µm.

**Supplementary Figure 2:**
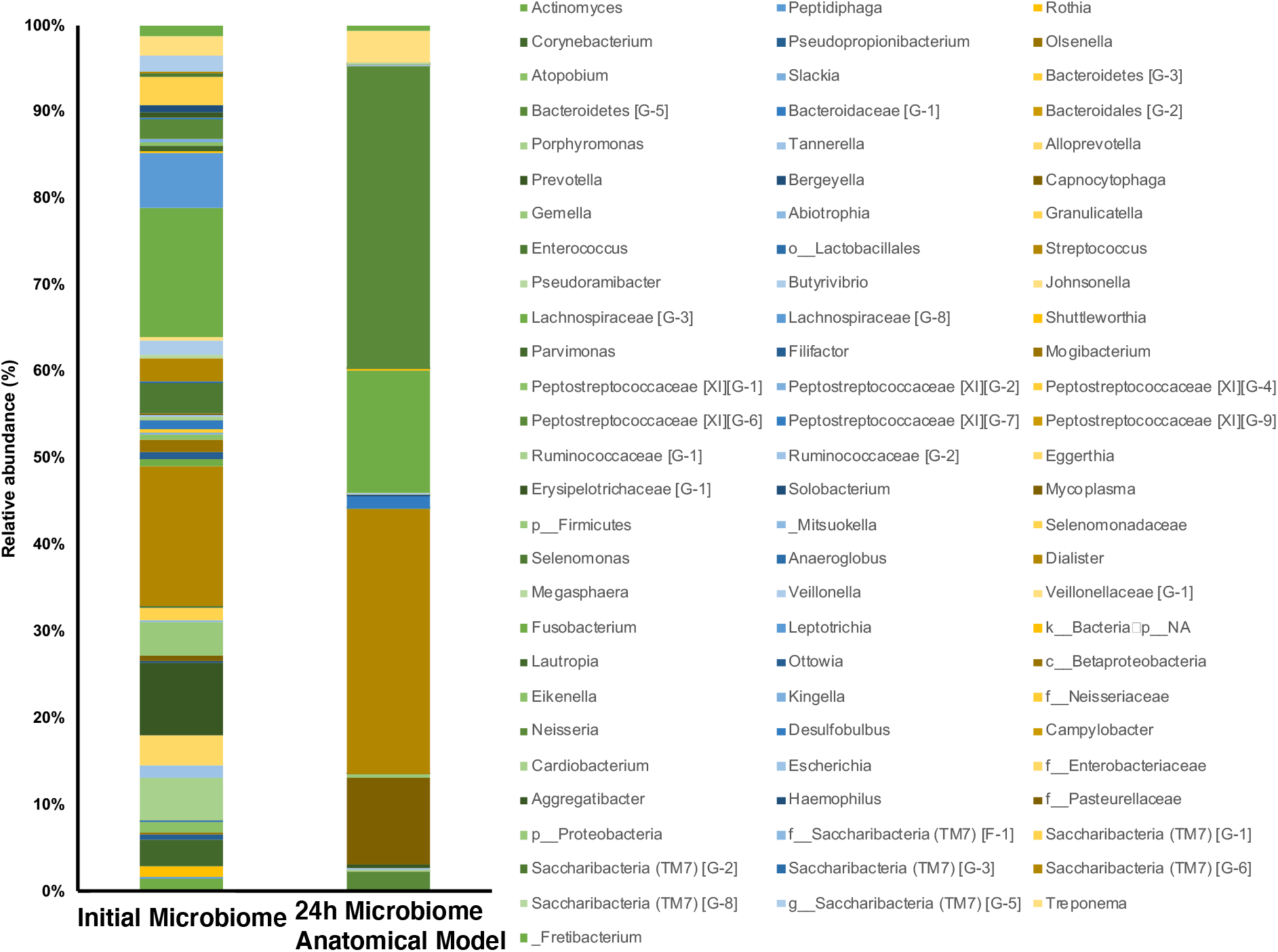
Relative abundance distribution of main genera detected in human plaque before incubation (Initial Microbiome) samples and samples collected after 1 day incubation in the anatomical model (24h Microbiome Anatomical Model).

**Supplementary Table 1:**
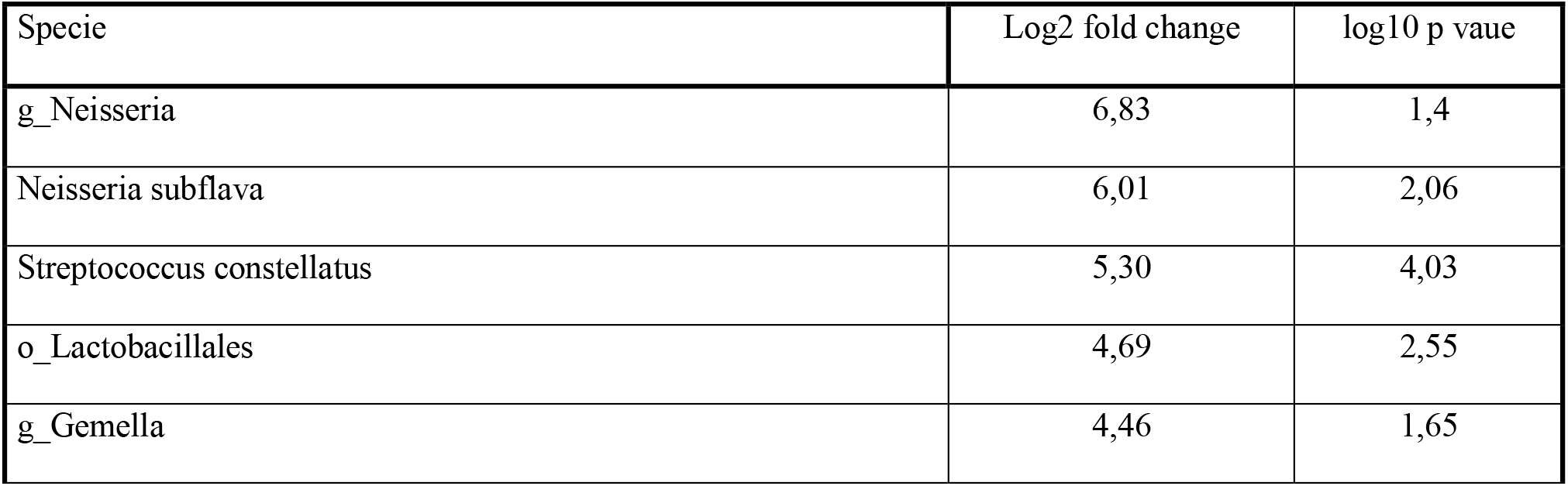
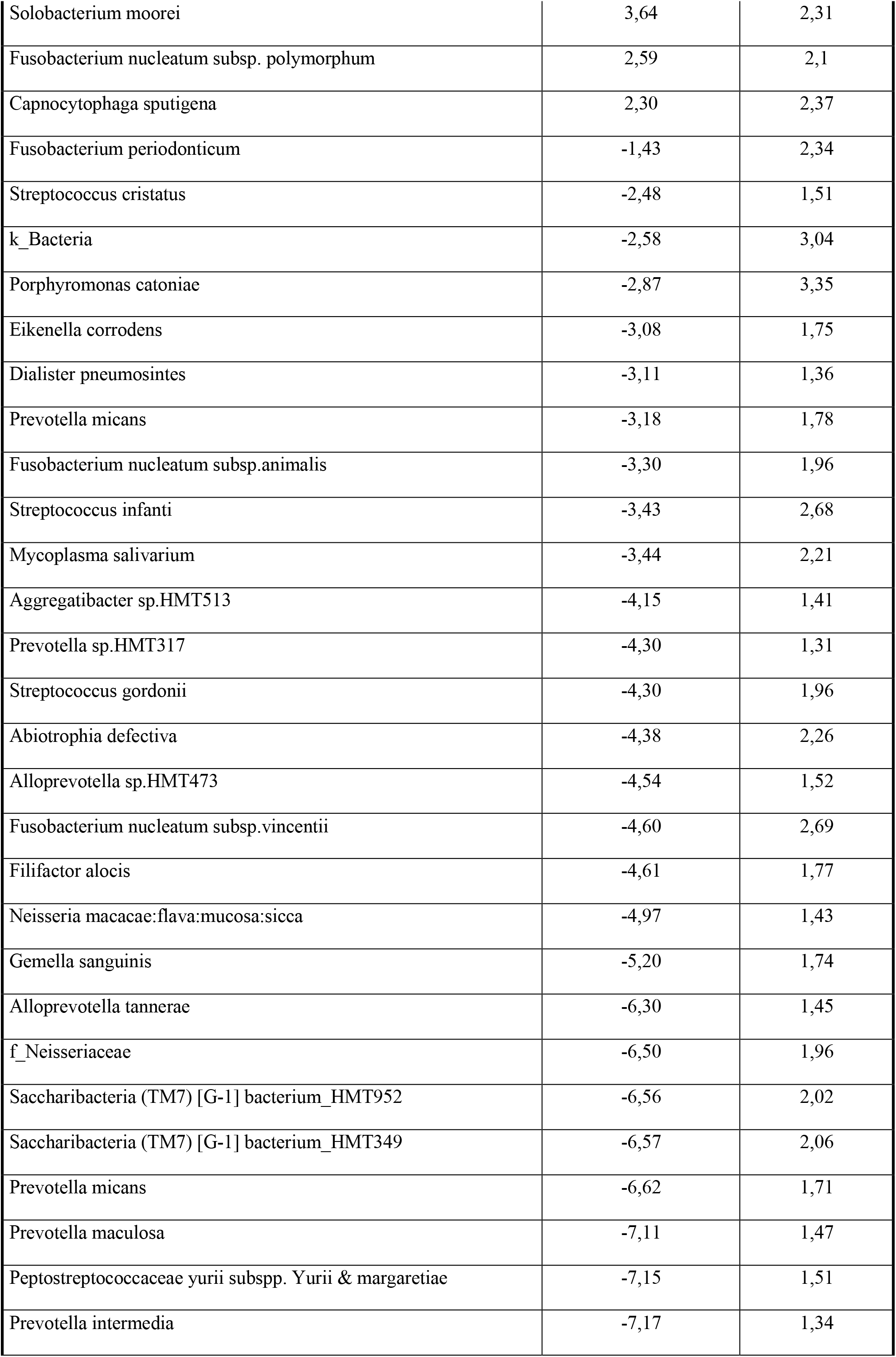
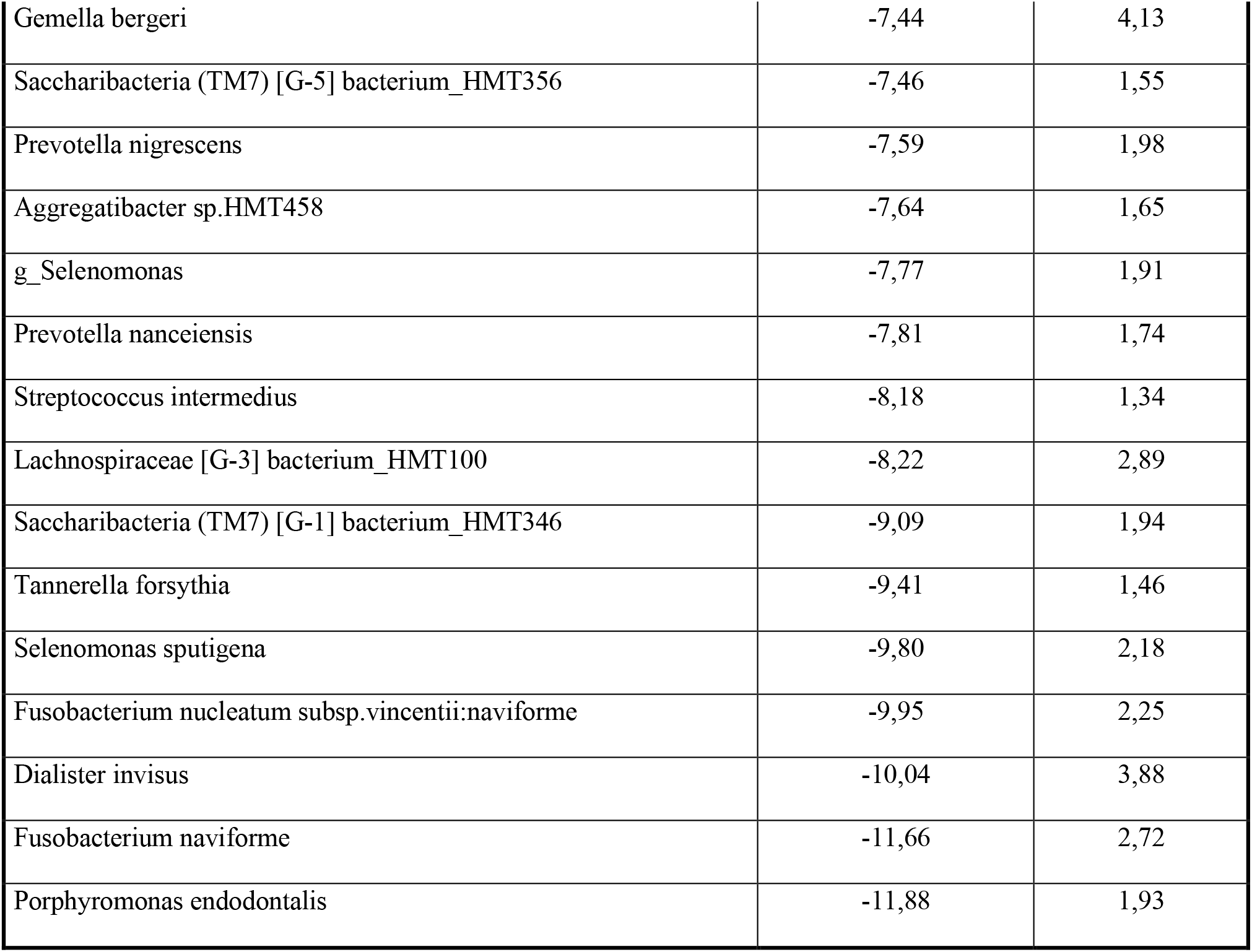
Statistical differences in changes of relative abundance at the level of specie. The table summarizes the Fold change and p-value with a 95% confidence.

